# Mechanical role of dural structures and tumor material heterogeneity in brain tumor mass effect

**DOI:** 10.64898/2026.11.26.769431

**Authors:** Nicole Tueni, Jonas Weidner, Marco Wittrich, Benedikt Wiestler, Silvia Budday

**Author notes:** corresponding author. *Email address:* (Silvia Budday).

## Abstract

Brain tumor growth induces considerable deformation of the surrounding tissue, yet the mechanical contribution of intracranial dural structures and the influence of tumor material heterogeneity on mass effect remain poorly understood. In this study, we developed a heterogeneous finite element model of the brain to investigate the mechanical role of the falx cerebri and tentorium cerebelli and to determine how variations in tumor subregion stiffness influence the mechanical response of the brain. Two complementary analyses were performed. First, synthetic lesion models with and without the falx and tentorium were compared under otherwise identical conditions. Inclusion of these structures produced only minor changes in global displacement magnitude, but substantially altered the spatial distribution of stress. The dural structures acted as stiff internal partitions that limited interhemispheric and supratentorial–infratentorial deformation and modified load transmission throughout the brain. Including the dural structures led to a different pattern of ventricular volume change during tumor growth, with both lateral ventricles compressed in the dura-included model, whereas without the dura the ventricle on the tumor side expanded while the opposite ventricle was compressed. Second, a parametric analysis was performed using MRI-informed patient-specific tumors comprising necrotic tumor core (NTC), contrast-enhancing tumor core (ETC), and peritumoral edema (Ed). Variations in ETC stiffness produced the largest and most spatially distributed changes in displacement and von Mises stress, with stronger effects in ETC-dominant tumors. Edema stiffness produced weaker and more diffuse effects, whereas NTC stiffness mainly affected the principal stretch within the tumor.

These findings demonstrate that intracranial dural structures alter the distribution of tumor-induced deformation and stress despite their limited influence on global displacement magnitude. In the patient-specific models, varying the tumor stiffness within its reported range changed the mechanical response by only a few percent, with the ETC having the largest influence.

## 1. Introduction

Tumor growth is a complex, multiscale process governed by the interplay of biological, biochemical, and mechanical factors. At the cellular and tissue scales, biological processes, such as cell proliferation, apoptosis, angiogenesis, hypoxia, and necrosis regulate tumor expansion, while biochemical mechanisms control nutrient transport and consumption [1, 2, 3]. Beyond these processes, mechanical interactions between the growing tumor and the surrounding host tissue play a critical role in shaping tumor morphology, redistributing stresses, and inducing deformation of adjacent healthy structures [4, 5]. Moreover, studies have demonstrated that solid stress accumulation within tumors can compress blood vessels, reduce perfusion, and significantly influence tumor progression [6]. Recent studies have highlighted the importance of mechanical coupling between tumor and host tissue in determining stress transmission and deformation [7]. These findings underscore the need to explicitly understand mechanical interactions when studying tumor growth.

In the brain, the mechanical effects associated with tumor growth are particularly significant due to the confined intracranial environment imposed by the skull. As tumors expand, they exert forces on surrounding brain tissue, leading to clinically relevant mass effect phenomena, such as midline shift, ventricular compression, and elevated intracranial pressure. These effects strongly influence the severity of neurological symptoms and the patient’s prognosis [8, 9].

Computational modeling of glioma growth has mainly focused on diffusion–reaction models that combine tumor cell proliferation with spatial invasion [2, 10]. Swanson’s Fisher-type formulation has provided the basis for many subsequent approaches, including extensions incorporating anisotropic diffusion to better capture invasion patterns across gray and white matter [2, 11]. Image-guided frameworks further allowed the estimation of patient-specific parameters from imaging data [11]. More recent developments included heterogeneous reaction–diffusion formulations, coupled with deformable image registration, and deep-learning surrogates on brain meshes [12]. In parallel, biomechanical “mass effect” models have been developed to simulate tumor-induced brain deformation. Early work relied on statistical or nonlinear elastic registration approaches combined with biomechanical formulations to estimate tumor-driven displacement fields, without explicitly modeling tumor growth dynamics [13, 14]. Subsequent finite element models reproduced observed displacements by prescribing growth or pressure-like loads within brain geometries, treating the tumor as a mechanical inclusion rather than a proliferating one [15, 16, 17]. More recent studies have examined the mechanical impact of tumor expansion on surrounding tissue and vasculature [18]. In these approaches, tumor growth is not modeled explicitly, and deformation arises solely from imposed mechanical loading.

More recently, inversion frameworks have been developed to extract physics-based biomarkers of mass effect directly from single-timepoint clinical imaging, using biomechanical growth models to infer patient-specific tumor and tissue parameters from the observed deformation field [19].

More advanced approaches couple reaction–diffusion models with biomechanical deformation, enabling simultaneous simulation of tumor cell-density evolution and mass effect. Such models have been used to generate realistic synthetic tumor datasets in MR images and to investigate how deformation influences the apparent morphology and extent of gliomas [20, 10]. Image-driven frameworks have further leveraged these models to estimate tumor growth parameters from patient data, linking observed deformations to underlying tumor cell distributions [21, 22].

Despite the advancements these frameworks provide, most existing biomechanical brain tumor models still rely on simplified tumor geometries and assume homogeneous or only weakly heterogeneous material properties for the brain. Recent experimental and computational studies have emphasized important spatial variability in mechanical properties across brain regions, as well as the importance of incorporating anatomically informed heterogeneity for accurate prediction of strain and stress fields in full-scale brain simulations [23, 24, 25].

In addition, the dura mater, including the falx cerebri and tentorium cerebelli, forms a complex, stiff partition that mechanically separates the major regions of the brain. Its role in the regulation of tumor-immune interactions has been increasingly studied, with recent work in glioblastoma and cerebral metastases, highlighting how dural immune cells and secreted factors can influence tumor growth and invasion [26, 27, 28, 29]. However, from a purely mechanical “mass-effect” perspective, the contribution of falx and tentorium to the redistribution of intracranial stresses and stretches under tumor-induced deformation has not yet been investigated to our knowledge. Moreover, ventricular displacement is a clinically established marker of mass effect [30]. Computational efforts to quantify the associated volumetric compression remain scarce. Ballatore et al. proposed a multiphase poroelastic model of anisotropic tumor growth reporting a 12.4% reduction in lateral ventricle volume over 140 days of simulated growth [31], while Subramanian et al. incorporated mass effect into an inversion-based frame-work to estimate tumor growth parameters from clinical imaging [19]. However, neither framework has addressed whether falx cerebri and tentorium cerebelli modulate the evolution of ventricular compression.

The mechanical properties of brain tumor tissue remain incompletely characterized and exhibit substantial variability across patients, tumor grades, and intratumoral regions. Reported stiffness values depend strongly on measurement modality, loading conditions, and whether measurements are performed *in vivo* or *ex vivo*, resulting in a wide range of estimates in the literature. Consequently, it remains unclear to what extent uncertainty in tumor mechanical properties affects predictions of tumorinduced brain deformation, stress redistribution, and clinically relevant measures of mass effect.

In this work, we deliberately neglect the biochemical aspects of tumor growth and focus exclusively on the mechanical interaction between the expanding tumor and the surrounding brain tissue. We address two main questions. First, we investigate how the inclusion of the falx cerebri and tentorium cerebelli influences tumor-induced deformation, stress and stretch redistribution, and ventricular compression, thereby assessing whether these dural structures must be represented explicitly in biomechanical models of mass effect. Second, we investigate the sensitivity of the predicted mechanical response to variations in tumor-region material properties.

To this end, we construct a heterogeneous finite element model of the brain comprising nine mechanically distinct anatomical regions and explicitly incorporate the falx cerebri and tentorium cerebelli. The framework is first applied to synthetic tumor configurations to isolate the mechanical influence of these dural structures on tumor-induced mass effect. Subsequently, we apply the model to MRI-informed patient-specific tumor geometries and systematically vary tumor material properties to quantify their influence on the global mechanical response.

## 2. Methods

This study consisted of two complementary mechanical analyses.

First, a synthetic lesion model was used to isolate the influence of the intracranial dural partitions, specifically the falx cerebri and tentorium cerebelli, on tumor-induced brain deformation. Two otherwise identical model configurations were compared: one including the falx and tentorium and one excluding these structures. All remaining model parameters, growth conditions, and boundary conditions were kept unchanged between the paired simulations.

Second, patient-specific tumor models were generated from preoperative MRI data to assess the sensitivity of the predicted brain mechanical response to tumor-subregion stiffness. Tumors were represented as heterogeneous structures composed of necrotic tumor core (NTC), contrast-enhancing tumor core (ETC), and peritumoral edema (Ed). The shear modulus of each tumor subregion was varied independently while the geometry, prescribed growth, surrounding tissue properties, boundary conditions, and remaining tumor-region parameters were held fixed. Changes in displacement, stretch, and stress fields were then quantified.

### 2.1. 3D brain models

#### 2.1.1. MRI processing, segmentation, and mesh generation Two distinct datasets were used to generate the finite element models

For the synthetic lesion analysis, structural brain MRI data were obtained from the OASIS (Open Access Series of Imaging Studies) cross-sectional dataset^1^. Anatomical brain segmentation was performed using the FreeSurfer image analysis suite [32], using the automated recon-all pipeline, which provides cortical and subcortical anatomical labels. The resulting segmentation masks were converted into volumetric meshes, and synthetic tumor lesions and dural structures were incorporated, as shown in Figure 1. A synthetic lesion configuration was adopted to allow controlled tumor size and placement. The lesion was positioned within the supratentorial white matter, lateral to the lateral ventricles and relatively distant from the tentorium cerebelli. This setup enabled the evaluation of the influence of the falx cerebri and tentorium cerebelli on both the mechanical response and the ventricular volumetric response. An additional synthetic configuration was investigated in which the tumor was placed in the inferior supratentorial region, immediately adjacent to the tentorium cerebelli, as shown in Figure C.15. The resulting deformation and stress patterns were qualitatively consistent with those obtained for the tumor positioned lateral to the lateral ventricles. The corresponding displacement and von Mises stress fields are shown in Figures C.16 and C.17, respectively, in Appendix C.

**Figure 1.**
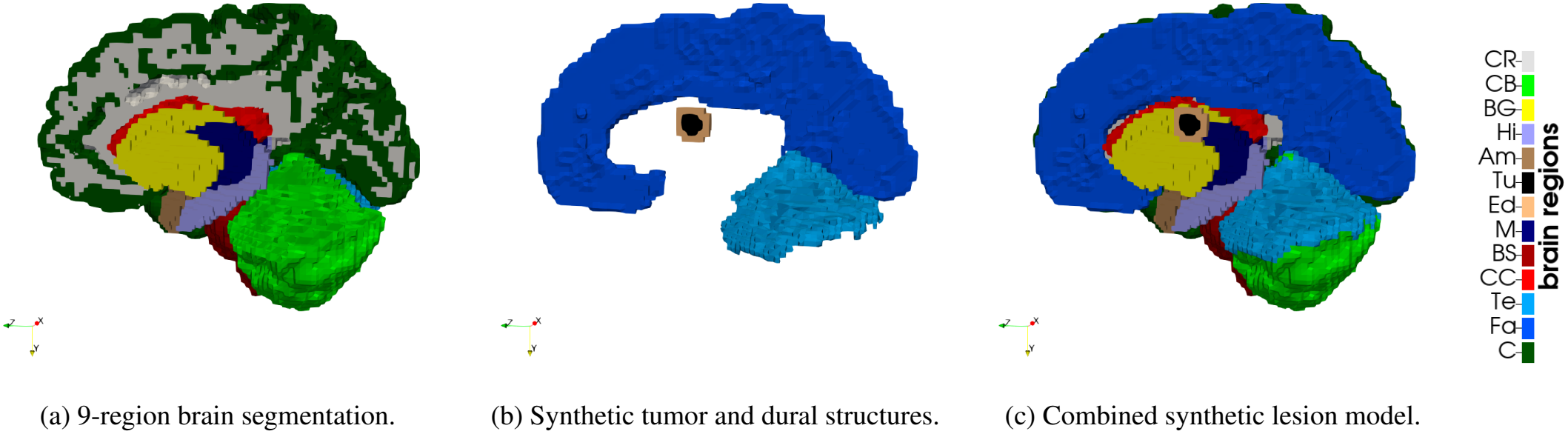
Finite element model used for the synthetic lesion study. The model includes nine mechanically distinct brain regions, dural structures represented by the falx cerebri and tentorium cerebelli, and a synthetic tumor with surrounding edema, placed lateral to the lateral ventricles.

For the patient-specific tumor analysis, preoperative MRI data and tumor segmentations were obtained from the BraTS 2021 dataset^2^. Tumor labels corresponding to necrotic tumor core, contrast-enhancing tumor core, and peritumoral edema were directly obtained from the dataset. Anatomical brain segmentation was again performed using FreeSurfer. The resulting anatomical brain masks were then merged with the BraTS tumor segmentation masks to generate a combined label map containing both healthy brain regions and tumor subregions. This final combined segmentation mask was subsequently meshed, and dural structures were added, as shown in Figure 2.

**Figure 2.**
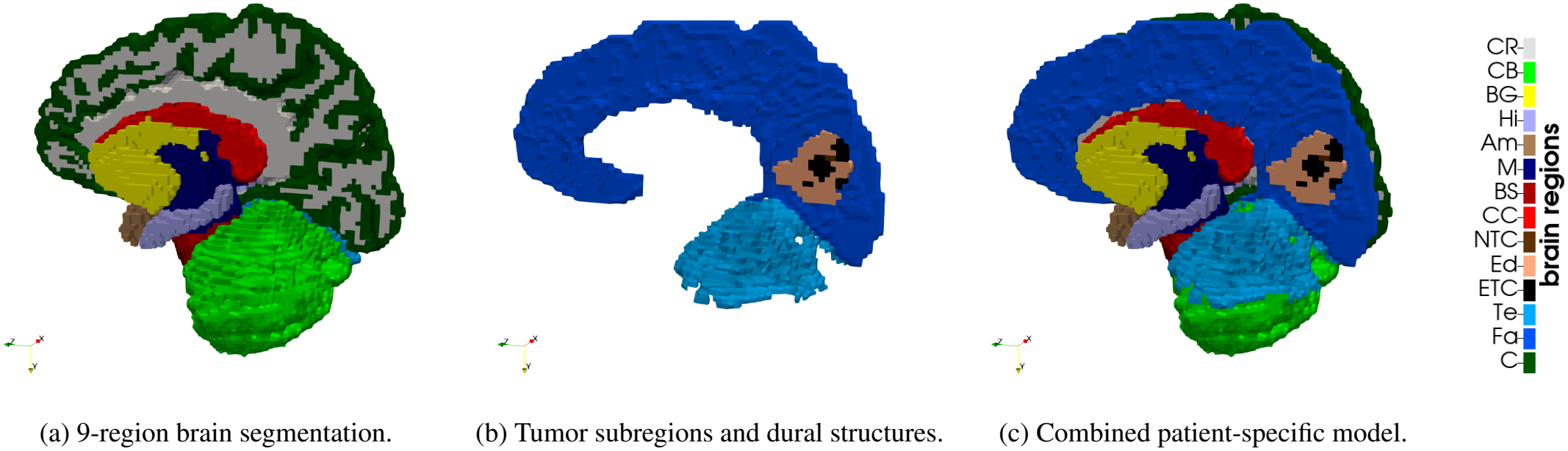
Patient-specific brain-tumor finite element model generated from patient MRI. The model combines the nine-region brain segmentation, falx and tentorium representation, and tumor subregions corresponding to necrotic tumor core, contrast-enhancing tumor core, and edema.

The segmented T1-weighted volumes were initially converted into structured hexahedral meshes with a spatial resolution of 1 × 1 × 1 mm^3^. To reduce computational cost, the meshes were coarsened to 2 × 2 × 2 mm^3^ elements while preserving the overall brain geometry. Mesh generation and coarsening were performed using an in-house Python-based meshing script^3^. Previous analyses demonstrated that simulations on 1 × 1 × 1 mm^3^ and 2 × 2 × 2 mm^3^ meshes produced comparable results [25].

Finally, Laplacian surface smoothing was applied to anatomical interfaces, and region-specific information was extracted for material assignment.

#### 2.1.2. Dura mater: tentorium and falx

Since none of the available automated tools currently provides explicit segmentation of intracranial dural structures suitable for finite element model generation, dural structures were incorporated using an automated approach based on the method proposed by Glaister et al. [33]. This method was implemented in the in-house meshing script and used to represent the dura mater through two main structures: the tentorium cerebelli, modeled as a layer between the cerebellum and cerebrum, and the falx cerebri, modeled as a layer separating the cerebral hemispheres, as shown in Figure 1 (b). Both structures are one element thick, and their thickness of 2 mm is thus set by the mesh resolution. This exceeds the measured thickness of the human falx cerebri of 0.91 ± 0.05 mm [34] and the values typically used in computational head models, 1.5 mm [35]. The mechanical influence of the dural structures is therefore likely overestimated, and the effects reported below should be regarded as an upper bound. The falx and tentorium were embedded in the hexahedral mesh, shared nodes with the adjacent tissue, and were not directly attached to the skull boundary. In the dura-excluded configuration, their elements were assigned CSF properties, so that both configurations use the same mesh.

### 2.2. Constitutive model and material parameters

Since the time scale is on the order of days, we neglected viscous and porous effects and modeled brain tissue as an isotropic and hyperelastic solid, using a one-term Ogden model, which has been shown to represent the time-independent mechanical response of brain tissue under multiple loading modes [23, 36]. The strain energy density function is decomposed into an isochoric and a volumetric part [37],

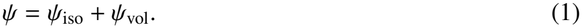

The isochoric part was defined in terms of the isochoric principal stretches 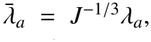, where *J* = det **F** and **F** is the deformation gradient,

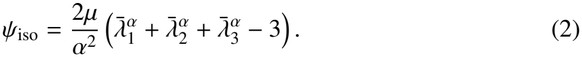

The volumetric part was defined as

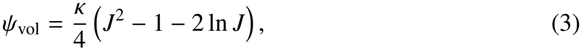

where *κ* is the bulk modulus, obtained from the shear modulus *µ* and Poisson’s ratio *ν* as

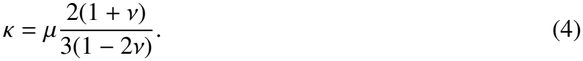

The Cauchy stress tensor was computed as

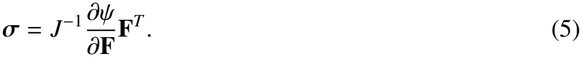

The brain was represented using the nine-region segmentation proposed by Hinrichsen et al. [38], consisting of cortex, corona radiata, basal ganglia, brainstem, midbrain, cerebellum, amygdala, hippocampus, and corpus callosum. This level of regional heterogeneity has previously been shown to capture important regional effects in brain tissue mechanics [24, 25]. A Poisson’s ratio of *ν* = 0.45 was assigned to all brain regions to represent slight compressibility of the brain tissue and the preconditioned state associated with slow tumor growth. Material parameters for each anatomical region are reported in Table A.3 in Appendix A. The cerebrospinal fluid (CSF) surrounding the brain and the ventricles were modeled as a very soft, compressible solid using the same one-term Ogden formulation, with a low stiffness *µ* = 10 Pa and *α* = −1.

The dura mater, falx cerebri, and tentorium cerebelli were modeled using the modified one-term Ogden parameters reported by De Kegel et al. [39], with *µ* = 0.47 MPa and *α* = 0.79, and a Poisson’s ratio of *ν* = 0.45.

#### 2.2.1. Synthetic tumor and edema material calibration

The synthetic lesion used to investigate the role of the dural structures was represented by two regions: a tumor core and a surrounding edema zone. Both regions were modeled using the same one-term Ogden hyperelastic formulation as the host tissue, with mechanical contrast introduced through variations in the shear modulus.

*In vivo* magnetic resonance elastography studies indicate that gliomas are generally softer than host white matter, whereas peritumoral edema has stiffness values comparable to, or only slightly lower than, healthy tissue [40, 41]. Relative to the white matter shear modulus used in this study, *µ*_WM_ = 157.09 Pa, the synthetic tumor was modeled as moderately softer than white matter, corresponding to a 10-20% reduction. The surrounding edema region was assigned a shear modulus up to 5% softer than the white matter. The adopted values were therefore

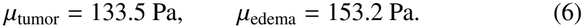

The Ogden nonlinearity parameter *α* was kept identical to that of the white matter host tissue, with *α* = −19.70.

#### 2.2.2. Patient-specific tumor subregion material calibration

For patient-specific simulations derived from MRI segmentation, the tumor was decomposed into three mechanically distinct subregions: necrotic tumor core, contrast-enhancing tumor core, and peritumoral edema. As in the synthetic study, all tumor regions were modeled using the same one-term Ogden hyperelastic formulation, with heterogeneity introduced through region-specific shear moduli.

MRE-based intratumoral analyses report relative stiffness contrasts within gliomas, with enhancing regions moderately softer than white matter and necrotic regions exhibiting the lowest effective stiffness [42]. Using the white matter reference value *µ*_WM_ = 157.09 Pa, the contrast-enhancing tumor core was modeled as 10-20% softer, while the necrotic tumor core was modeled as 20-30% softer. Thus,

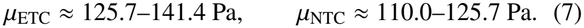

Following the synthetic study, the edema region was modeled with a maximum stiffness reduction of 5% compared to healthy white matter,

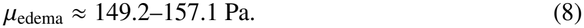

As before, the one-term Ogden nonlinearity parameter *α* was kept equal to that of the surrounding white matter.

### 2.3. Tumor growth formulation

Tumor growth was modeled using a multiplicative decomposition of the deformation gradient into an elastic part **F**^*e*^ and a growth part **F**^*g*^,

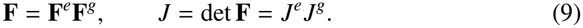

Growth was assumed to be isotropic,

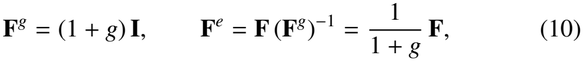

where *g* ≥ 0 is a scalar, dimensionless growth coefficient. In the synthetic lesion model, both the tumor core and the surrounding edema grew. In the patient-specific models, growth was restricted to the contrast-enhancing tumor core and the edema, and the necrotic tumor core was treated as a passive region. Healthy parenchyma, dural structures, CSF, and ventricles were assigned *g* = 0, so that **F**^*e*^ = **F** in all non-growing tissue. Tumor expansion and edema swelling therefore followed the same growth law, and the growing regions differed only in their material parameters.

The strain energy function, Equations 1–3, was evaluated with the elastic deformation gradient **F**^*e*^, and stresses were computed from Equation 5, using **F**^*e*^ instead of **F**.

With isotropic growth, the growth-induced volume ratio is

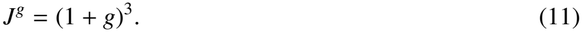

To relate the growth coefficient to clinically observed tumor volume doubling times (VDT), we assumed exponential volume growth of the growing regions, *J*^*g*^(*t*) = 2^*t/*VDT^, which gives

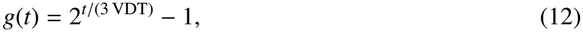

where *t* is the growth time in days. For small *t*, this corresponds to a growth rate of

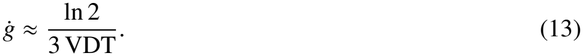

Reported preoperative glioblastoma doubling times of 20-50 days [43] thus translate into growth rates of roughly 0.005-0.012 per day. All simulations used VDT = 20 days.

For the synthetic lesion, the growth coefficient was increased in 60 equal increments of Δ*g* ≈ 0.0136 up to *g*_max_ = 6^1*/*3^ − 1 ≈ 0.817. The patient-specific tumors required a larger number of simulations, so growth was applied in 10 increments of 2 days up to *t* = 20 days, reaching *g*_max_ = 2^1*/*3^ − 1 ≈ 0.260.

### 2.4. Boundary conditions

Boundary conditions followed our previous study [25]. Zero-displacement Dirichlet constraints were imposed on the outer boundary of the cerebrospinal fluid space, corresponding to the inner skull surface.

### 2.5. Study objectives

#### 2.5.1. Influence of dural structures

The first simulation analysis evaluated the mechanical influence of the falx cerebri and tentorium cerebelli on tumorinduced brain deformation. Synthetic lesion simulations were performed using two paired configurations: one including the dura and one excluding it. Model outputs, including displacement magnitude and von Mises stresses, were then compared between the two configurations.

In addition to global displacement and stress, the volumetric evolution of the ventricles was quantified over the course of simulated tumor growth, for both configurations. This analysis was motivated by clinical evidence that ventricular displacement is a correlate of mass effect and prognosis that is not fully explained by tumor volume [30], and by the absence of dural structures in existing computational models of ventricular compression and mass effect [31, 19].

Ventricular volume was computed as the summed volume of the deformed ventricle elements. To distinguish the two lateral ventricles, ventricle elements were assigned to the tumor side or the opposite side according to their position relative to the midline, defined by the falx, in the undeformed configuration. For each lateral ventricle, the lateral displacement of its center was additionally computed, following the definition of lateral ventricle displacement in [30].

#### 2.5.2. Influence of tumor subregion stiffness

The second simulation analysis evaluated the effect of tumorregion material parameters on brain mechanical response, using patient-specific brain-tumor models containing three tumor subregions: necrotic tumor core (NTC), contrast-enhancing tumor core (ETC), and peritumoral edema (Ed).

To separate material effects from volumetric effects, one control case was selected in which the three tumor subregions had approximately balanced volume fractions. Additional patientspecific cases with different tumor-subregion volume fractions were also analyzed to assess the influence of diverse tumor compositions.

A parametric analysis was performed to evaluate how tumor stiffness affects mechanical behavior. For each subregion, three shear modulus values were defined: a lower bound, a middle value, and an upper bound, based on the realistic ranges established above. The middle values were chosen as the baseline configuration. Each parameter was then varied independently to its lower and upper bounds, while the other two parameters were kept constant at their baseline values. This resulted in seven simulations: one baseline case and two variations per tumor region, as summarized in Table 1.

**Table 1.** Parametric analysis of tumor-subregion shear moduli with dural structures included. Lower-bound and upper-bound values of each tumor-subregion shear modulus were evaluated independently while the remaining parameters were maintained at their baseline values. Gray-shaded cells indicate shear moduli that were held fixed at the baseline values in each simulation, whereas unshaded cells denote the parameter being varied.

| Configuration | *μ*<sub>NTC</sub> (Pa) | *μ*<sub>ETC</sub> (Pa) | *μ*<sub>edema</sub> (Pa) |
|---|---|---|---|
| Baseline | 117.8 | 133.5 | 153.2 |
| Lower-bound NTC | 110.0 | 133.5 | 153.2 |
| Upper-bound NTC | 125.7 | 133.5 | 153.2 |
| Lower-bound ETC | 117.8 | 125.7 | 153.2 |
| Upper-bound ETC | 117.8 | 141.4 | 153.2 |
| Lower-bound edema | 117.8 | 133.5 | 149.2 |
| Upper-bound edema | 117.8 | 133.5 | 157.1 |

### 2.6. Numerical implementation

All simulations were performed with an in-house finite element code based on the open-source library deal.II ^4^ [44]. The synthetic lesion model comprised 175,145 elements and 561,150 degrees of freedom; the dura-excluded configuration used the same mesh. The patient-specific models comprised between 232,267 and 267,542 elements and between 739,782 and 849,717 degrees of freedom (Table A.4).

## 3. Results and Discussion

### 3.1. Effect of dural structures on tumor-induced brain deformation

#### 3.1.1. Displacement

Figure 3 shows the simulated displacement field for models that include and exclude the dura mater (columns 1 and 2 respectively), the absolute difference between the two configurations (column 3), the displacement field within the dura (column 4), and the anatomical regions for reference (column 5).

**Figure 3.**
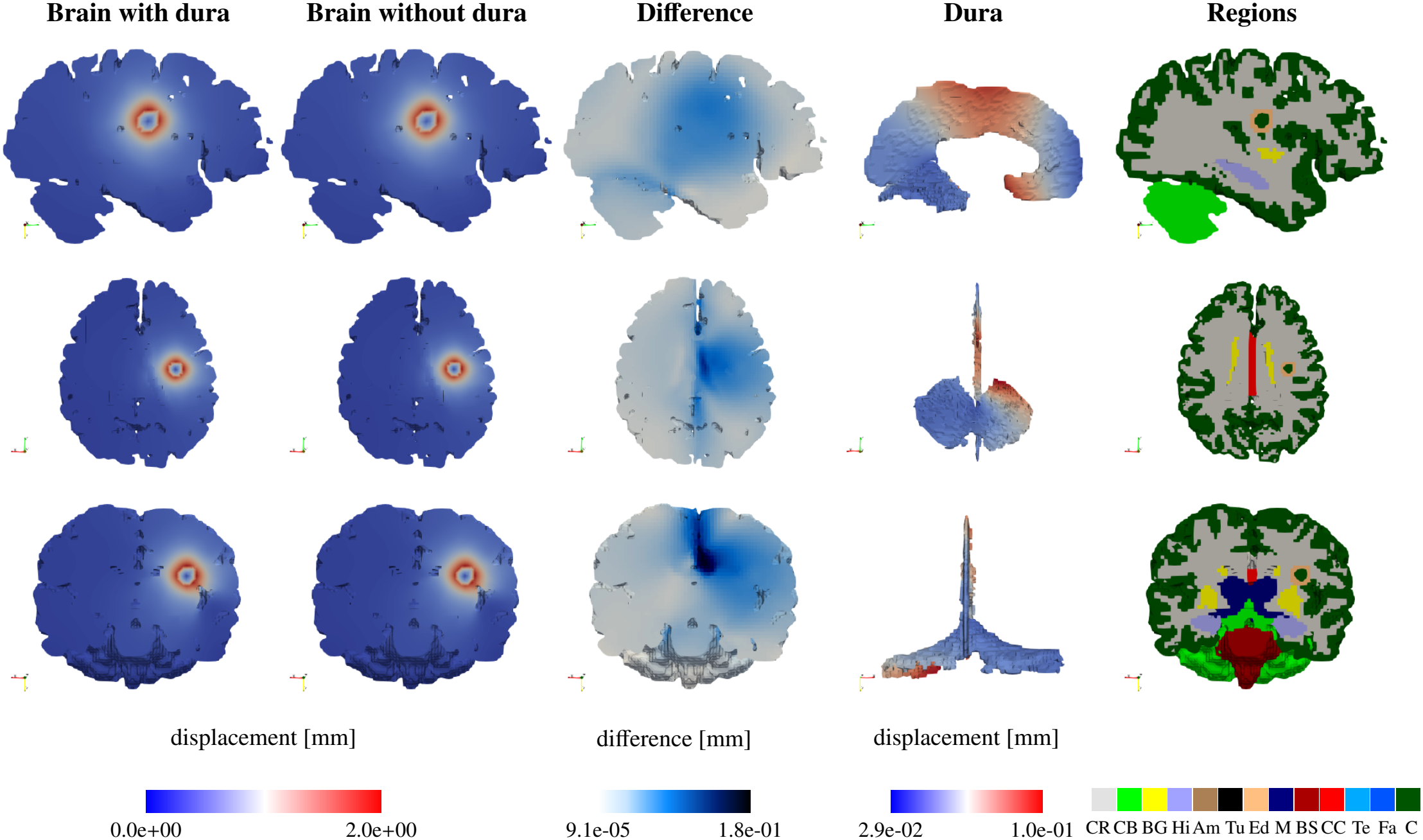
Displacement field [mm] comparison between models with and without dural structures for the synthetic lesion. Columns show displacement magnitude with dura, displacement magnitude without dura, absolute displacement difference, displacement of the dural structures, and anatomical regions. Rows correspond to sagittal, axial, and coronal sections.

Columns 1 and 2 demonstrate qualitatively similar global displacement patterns. In both configurations, the displacement field is strongly localized around the tumor, with a smoothly decreasing magnitude with distance from the lesion. The peak displacement reaches approximately 2.0 mm and remains centered at the tumor in both models. This reflects the dominant role of tumor volumetric growth combined with the confinement imposed by the Dirichlet boundary condition at the skull. The absolute displacement difference between the two configurations is comparatively small, with a maximum of approximately 0.18 mm, corresponding to about 9% of the peak displacement magnitude. These results indicate that inclusion of the falx and tentorium does not substantially alter the overall magnitude of tumor-induced mass effect.

However, despite its small magnitude, the displacement difference exhibits clear spatial organization rather than a uniform distribution. In the sagittal view, the tentorium cerebelli introduces a mechanical separation between supratentorial and infratentorial regions, mitigating inferior displacement of the cerebellum and brainstem. In axial and coronal views, large differences cluster along the interhemispheric region, consistent with the falx cerebri limiting interhemispheric sliding. These results were also reproduced with the supratentorial tumor, as observed in Figure C.16.

Column 4 shows that both falx and tentorium undergo small but non-negligible displacements. The displacement field is anisotropic and concentrated in regions near the tumor, reinforcing the interpretation of the dura as a stiff, load-bearing structural membrane rather than a passive boundary.

Overall, incorporating dural structures yields only minor changes in global displacement magnitude, but alters the spatial distribution of deformation throughout the brain. This redistribution may affect directional indicators of mass effect, although the absolute differences in the present simulations are small.

#### 3.1.2. Stresses

Figure 4 presents the corresponding von Mises stress fields for both configurations (columns 1 and 2), their absolute difference (column 3), the stress field within the dura (column 4), and the anatomical regions for reference (column 5).

**Figure 4.**
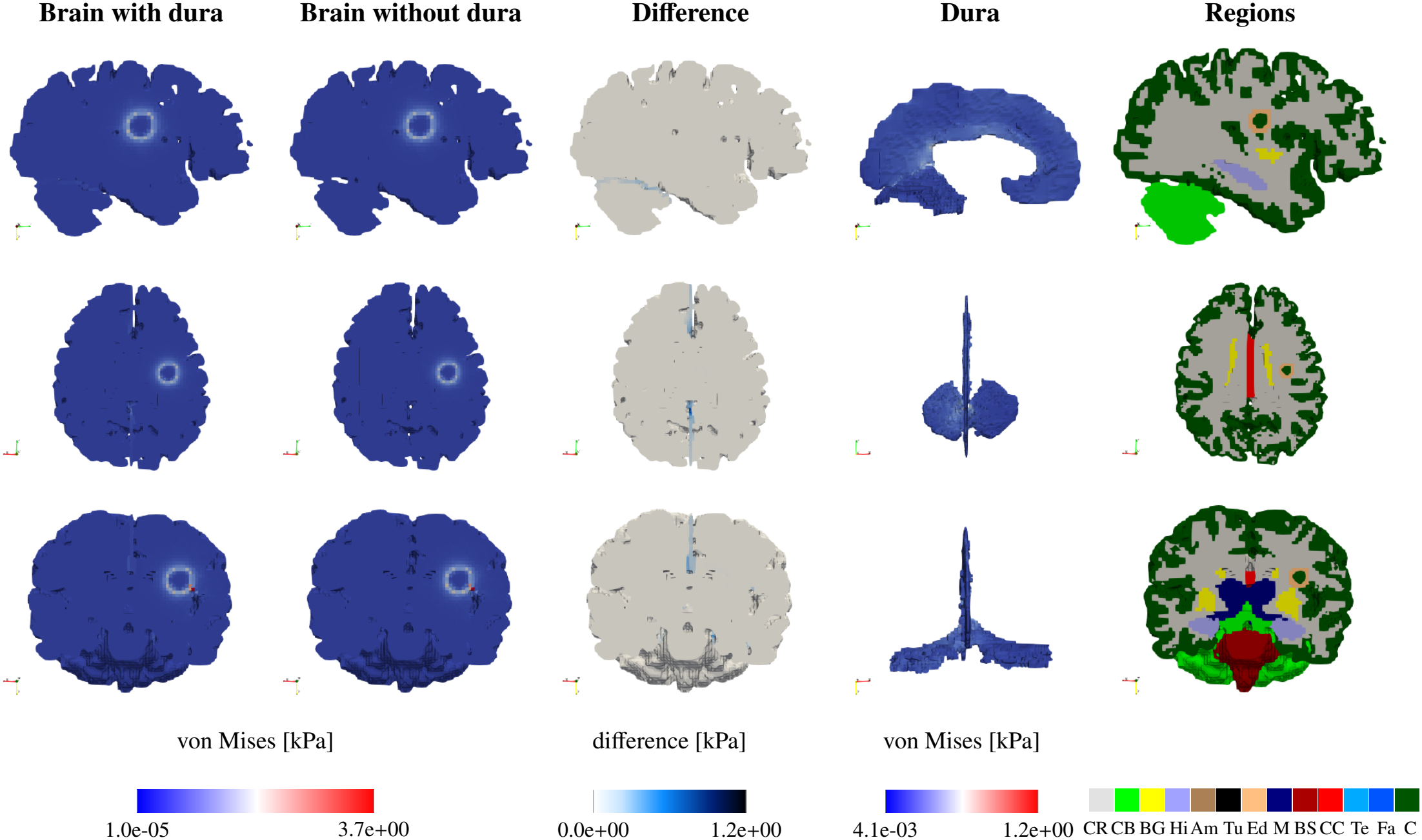
Von Mises stress field [kPa] comparison between models with and without explicit dural structures for the synthetic lesion. Columns show von Mises stress field with dura, von Mises stress field without dura, absolute stress difference, stress carried by the dural structures, and anatomical regions. Rows correspond to sagittal, axial, and coronal sections.

In both configurations, the highest von Mises stresses, reaching approximately 3.7 kPa, are localized around the tumor. These stress concentrations arise from growth-induced volumetric incompatibility between the expanding tumor and surrounding healthy host tissue, combined with global skull confinement. In contrast to the displacement field, von Mises stresses demonstrate greater sensitivity to the presence of the dura. The absolute stress difference reaches approximately 1.2 kPa, corresponding to about 32% of the peak stress magnitude.

The difference patterns in column 3 align strongly with dural structures, in all anatomical views, indicating that stress redistributes along these internal partitions. The dura modifies load transmission and alters stress localization at mechanical interfaces.

Column 4 shows that the dura itself carries non-negligible von Mises stress, with values up to approximately 1.2 kPa. Although its displacement remains small, the dura supports meaningful stress concentrations near the tumor.

These results, reproduced by the supratentorial tumor as observed in Figure C.17, indicate that the falx cerebri and tentorium cerebelli act as internal stiff membranes that function as load-bearing and separating interfaces between both cerebral hemispheres, as well as between the cerebrum and the cerebellum. Their influence is limited in terms of magnitude but more substantial in shaping deformation gradients and stress redistribution.

#### 3.1.3. Ventricular response

Figure 5a shows the evolution of lateral ventricle volume with increasing growth coefficient *g* for the lateral tumor configuration with and without dural structures. In the duraincluded configuration, ventricular volume decreased monotonically throughout the simulated growth range, reaching a total change of approximately −0.16%. In the dura-excluded configuration, ventricular volume decreased by only approximately 0.03% up to *g* ≈ 0.31 and then gradually returned toward its initial value, reaching −0.01% at the end of growth.

**Figure 5.**
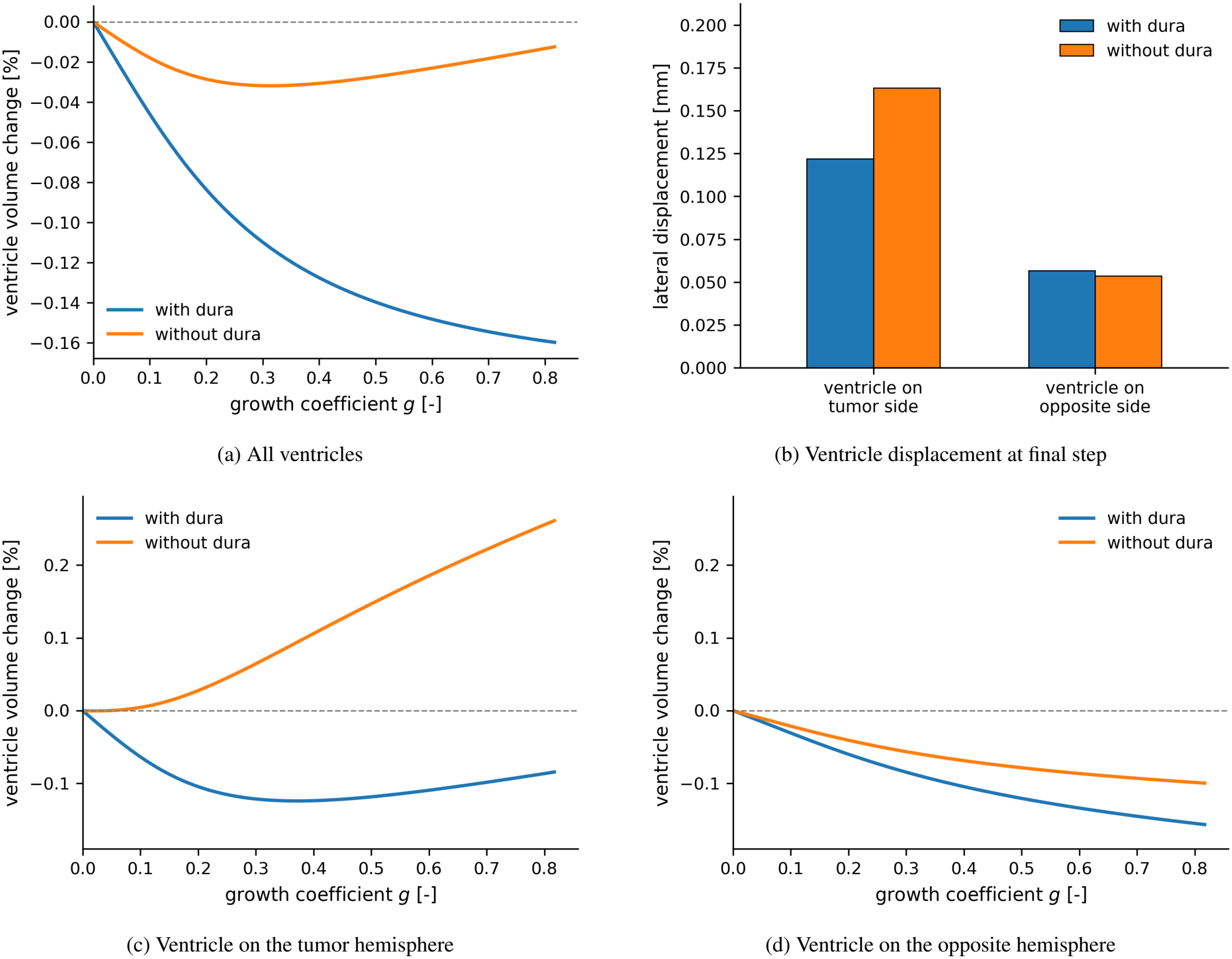
Ventricular response to tumor growth with (blue) and without (orange) dural structures, for the lateral tumor configuration. (a) Change in total lateral ventricle volume with increasing growth coefficient *g*. (b) Lateral displacement of the center of each lateral ventricle away from the tumor. (c, d) Volume change of the ventricle on the tumor hemisphere and of the ventricle on the opposite hemisphere.

To understand this difference, we computed the lateral displacement of the center of the ventricle on the tumor side and of the ventricle on the opposite side (Figure 5b). The lateral direction is the component captured by midline shift, and it dominates the displacement of the ventricles [30]. With the dura, the ventricle on the tumor side displaced less (0.122 mm versus 0.163 mm), whereas the ventricle on the opposite side displaced slightly more (0.057 mm versus 0.054 mm). The stiff falx thus acts as an internal wall that limits the displacement on the tumor side and transmits the load across the midline.

This load transfer is reflected in the volumes of the two lateral ventricles (Figures 5c and 5d). Without the dura, the two ventricles behaved in opposite ways. The ventricle on the tumor side, not constrained by the falx, expanded continuously during growth (approximately +0.26%), whereas the ventricle on the opposite side was compressed (approximately −0.10%), likely due to the confinement imposed by the Dirichlet boundary condition on the skull. The total ventricular volume therefore results from two opposing trends, which produce the partial recovery observed in Figure 5a. With the dura, the tissue surrounding the ventricle on the tumor side is confined between the growing tumor and the falx. This ventricle was therefore compressed by up to approximately 0.12% and recovered only partially, possibly because it expanded in the directions not constrained by the falx. At the same time, the load transmitted by the falx compressed the ventricle on the opposite side more strongly than without the falx (approximately −0.16%), so that the total volume decreased monotonically.

It should be emphasized that these observations are based on a mechanistic model of a single synthetic lesion, and the explanations given here are interpretations of the simulated deformation. The predicted changes are small and depend on model assumptions. Clinical validation, for example with longitudinal imaging of patients with laterally located tumors, is therefore required.

### 3.2. Patient-specific analysis of tumor material sensitivity

Based on the results of the above analysis, all subsequent patient-specific simulations included the falx cerebri and tentorium cerebelli. The analysis was then extended from synthetic lesions to MRI-derived patient-specific tumors. Multiple patient-specific brain-tumor models were segmented and meshed: one control case with a balanced volume fraction of tumor sub-regions, which serves as a baseline to decouple parameter effects from compositional variability, and three representative patients with different tumor compositions.

Figure 6 shows the evolution of tumor expansion with increasing values of the prescribed growth parameter *g* for the control and one representative patient-specific tumor.

**Figure 6.**
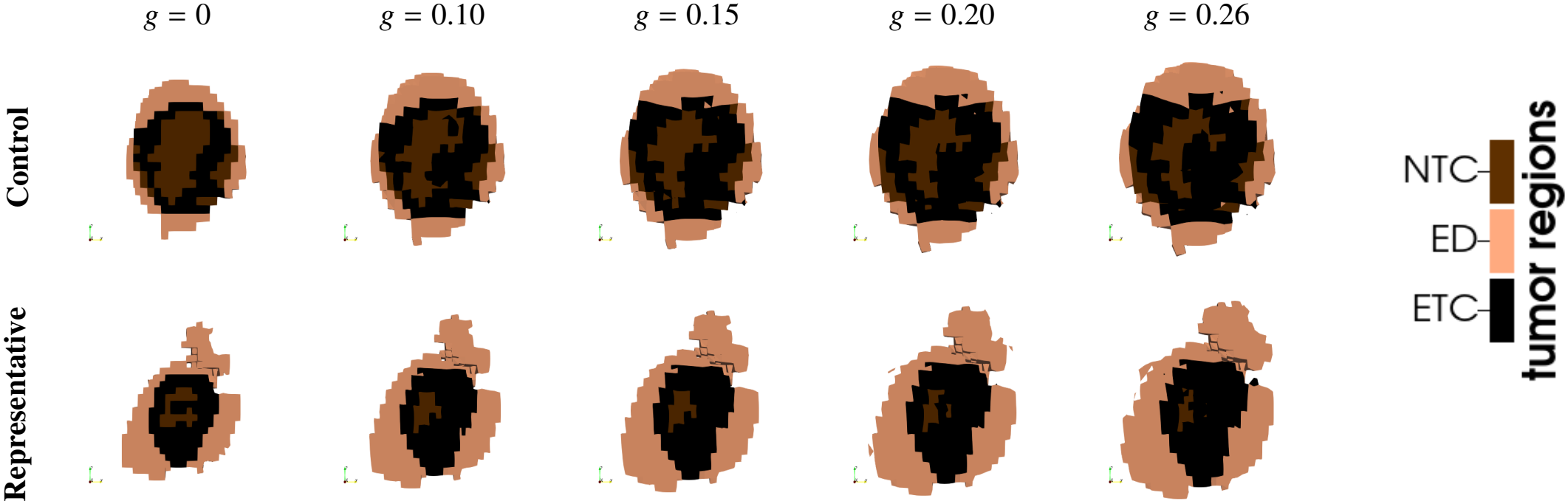
Tumor growth progression in the control and one representative patient-specific tumor for increasing values of the prescribed growth parameter *g*. The control case has approximately balanced tumor subregion fractions, whereas the representative case illustrates a heterogeneous patient-specific tumor morphology.

#### 3.2.1. Parametric analysis of tumor material parameters

For each brain configuration, simulations with baseline material parameters, as well as six additional cases with varied tumor material properties, as described in Section 2.5, were performed. This resulted in a total of seven simulations per brain. Each simulation was computed over multiple time steps corresponding to increasing values of the growth parameter *g*. Sensitivities were evaluated at the final growth step (*g* = 0.26).

*Control case*.

##### Displacement

Figure 7 presents, for the control case, the percentage change in the magnitude of maximum displacement (calculated as the regional p95 value) for each healthy brain region and tumor subregion, relative to the baseline simulation. This allows us to assess how localized variations in the properties of the tumor material influence both the tumor itself and the surrounding brain tissue. A clear hierarchical response was observed across the three perturbed tumor regions: variation of the ETC shear modulus produced the largest effect on displacement, followed by edema, whereas perturbations of the NTC produced only minor changes.

**Figure 7.**
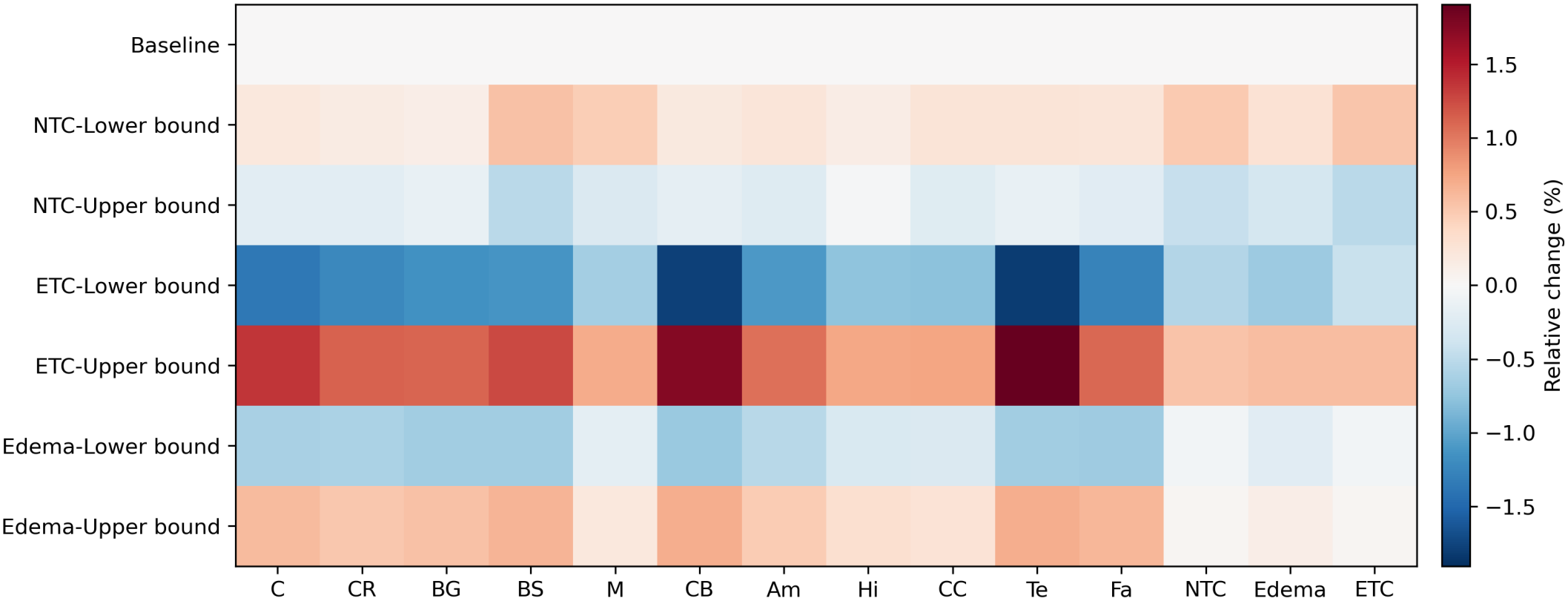
Control case sensitivity of regional displacement magnitude to tumor subregion stiffness perturbations. Values indicate percentage change relative to the baseline simulation.

The strongest displacement sensitivity was associated with ETC perturbations. Lowering *µ*_*ETC*_ produced a reduction in regional displacement across all regions, with the largest decreases observed in the cerebellum and dura mater, while increasing *µ*_*ETC*_ led to a corresponding increase (reaching up to approximately 1.5% variation). Edema perturbations produced the same trend, but with smaller amplitudes. Reducing edema stiffness resulted in a mild decrease in displacement (below 1%), whereas increasing it yielded a moderate increase, again across most healthy and tumor regions, and more specifically within the cerebellum and dura mater. Compared with the ETC, the edema effect was more diffuse across brain regions, which indicates a more buffering and spreading role, as it is the outer tumor-associated region directly in contact with several regions of the host tissue.

By contrast, changes in NTC stiffness had only a weak influence on displacement, with regional variations remaining small relative to those induced by ETC or edema perturbations. Note that decreasing *µ*_*NTC*_ led to an increase in displacement across nearly all regions relative to baseline, whereas increasing *µ*_*NTC*_ produced the opposite effect. Since the NTC does not undergo growth, it acts as a passive mechanical inclusion within the tumor mass: a softer NTC therefore provides less resistance to deformation induced by the surrounding growing ETC and edema, allowing slightly larger displacement, while a stiffer NTC mechanically resists tumor expansion and reduces the resulting displacement field.

Overall, some regions were more sensitive than others. The dura mater, cerebellum, and cortex exhibit large percentage changes in displacement, whereas the midbrain, hippocampus, and corpus callosum show only weak sensitivity across all parameter perturbations. This suggests that the mechanical response is not determined solely by the altered tumor material parameter itself, but is also influenced by the anatomical location of the tumor within the brain and by the constitutive properties of the surrounding healthy regions.

##### von Mises stress

Figure 8 shows the percentage change in regional von Mises stress relative to the baseline simulation for the control case. Overall, a sensitivity pattern similar to the displacement one is observed. Variations in the shear modulus of the ETC produce the largest changes in stress, followed by edema, while perturbations of the NTC result in smaller effects. The most pronounced response is associated with ETC perturbations. Increasing *µ*_*ETC*_ produces an increase in stress in all regions, with the largest augmentation occurring within the tumor itself, especially in the NTC and ETC regions, followed by the tentorium. Note that the stress increase within the passive NTC is particularly high. This can be explained by the fact that the growing contrast-enhancing tumor core leads to stress build-up in the neighboring passive necrotic tumor core. Decreasing *µ*_*ETC*_ leads to a reduction in von Mises stress across all regions, with strong decreases observed in the ETC itself, the tentorium, and the corpus callosum.

**Figure 8.**
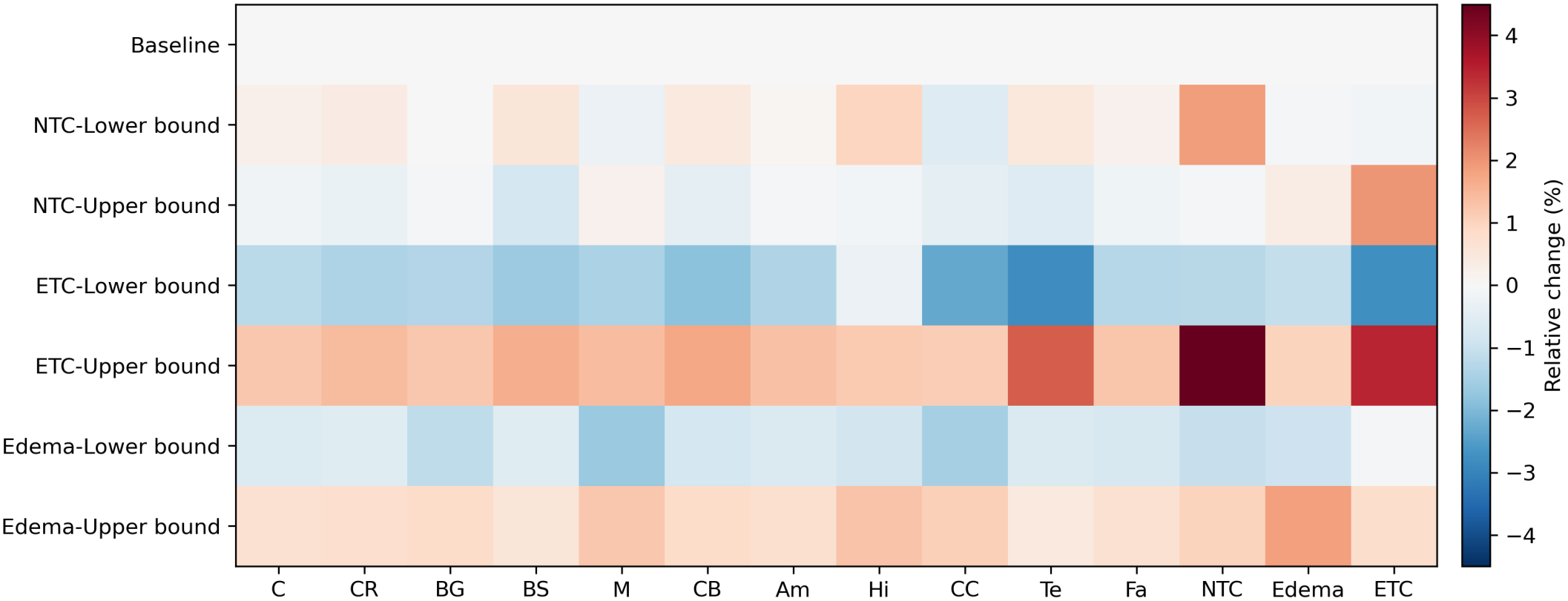
Control case sensitivity of regional von Mises stress to tumor subregion stiffness perturbations. Values indicate percentage change relative to the baseline simulation.

Edema perturbations produce a similar trend. Lowering *µ*_*Edema*_ leads to a moderate decrease in von Mises stress across most regions, while increasing it results in a corresponding increase. Compared with ETC, the effect of edema is more spatially uniform and less localized, with relatively similar changes observed across both tumor and healthy regions.

In contrast, variations in NTC stiffness produce a different sensitivity pattern in the von Mises stress field. Decreasing *µ*_*NTC*_ leads to a slight increase in von Mises stress within the NTC itself as well as across most surrounding brain regions. In contrast, increasing *µ*_*NTC*_ produces higher stress levels within the ETC and edema regions, while slightly reducing stress in most healthy tissues. Since the NTC is not subjected to growth, these trends suggest that its stiffness primarily influences how stresses are redistributed within the tumor mass. A softer NTC allows for more internal deformation and leads to a more diffuse stress distribution into the surrounding regions.

##### First principal stretches

Figure 9 shows the percentage change in the first principal stretch relative to the baseline simulation for the control case. Overall, the largest sensitivity is associated with perturbations of the NTC stiffness, followed by ETC, while edema perturbations produce comparatively small variations.

**Figure 9.**
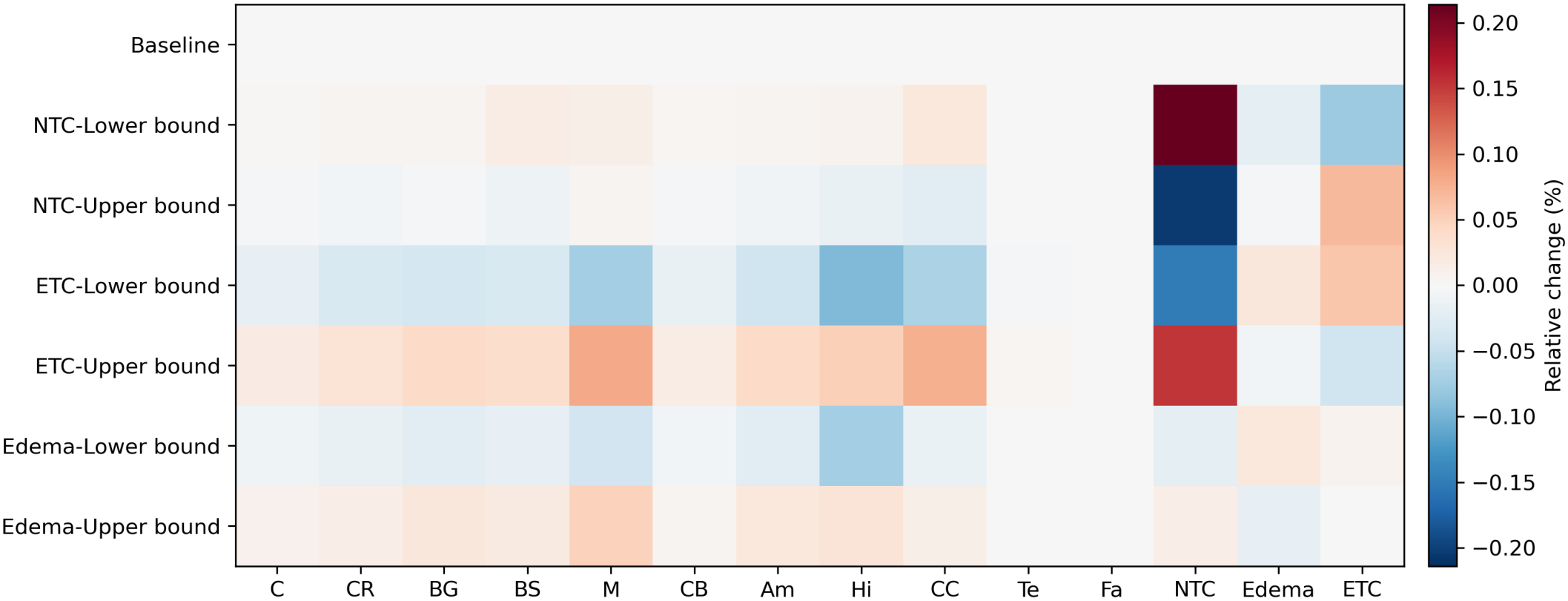
Control case sensitivity of regional first principal stretch to tumor subregion stiffness perturbations. Values indicate percentage change relative to the baseline simulation.

Decreasing *µ*_*NTC*_ produces the largest increase in first principal stretch within the NTC itself, as well as smaller increases across most healthy brain regions, while simultaneously leading to a decrease in the ETC region. The opposite behavior is observed when increasing *µ*_*NTC*_, with the strongest reduction in tensile stretch occurring within the NTC itself, together with mild decreases across most healthy tissues, while the ETC region exhibits the largest increase. Since the NTC is a passive region, these results indicate that a softer NTC allows larger tensile deformation within the necrotic core, whereas a stiffer NTC shifts the deformation to the surrounding ETC region.

This is followed by ETC perturbations. Decreasing *µ*_*ETC*_ leads to a reduction in the first principal stretch within the NTC region and across most healthy brain regions, while simultaneously increasing tensile stretch within the edema and ETC regions. In contrast, increasing *µ*_*ETC*_ produces a greater tensile stretch in the NTC region and surrounding healthy tissues, together with a reduced stretch within the ETC region.

Edema perturbations follow qualitatively similar trends to ETC perturbations but with smaller amplitudes. Increasing and decreasing *µ*_*Edema*_ produce only weak changes in the first principal stretch field.

##### Third principal stretches

Figure 10 shows the percentage change in the third principal stretch relative to the baseline simulation for the control case. In contrast to the first principal stretch, the sensitivity pattern is much more localized and concentrated within the tumor regions, with only minimal influence on the surrounding healthy brain tissue.

**Figure 10.**
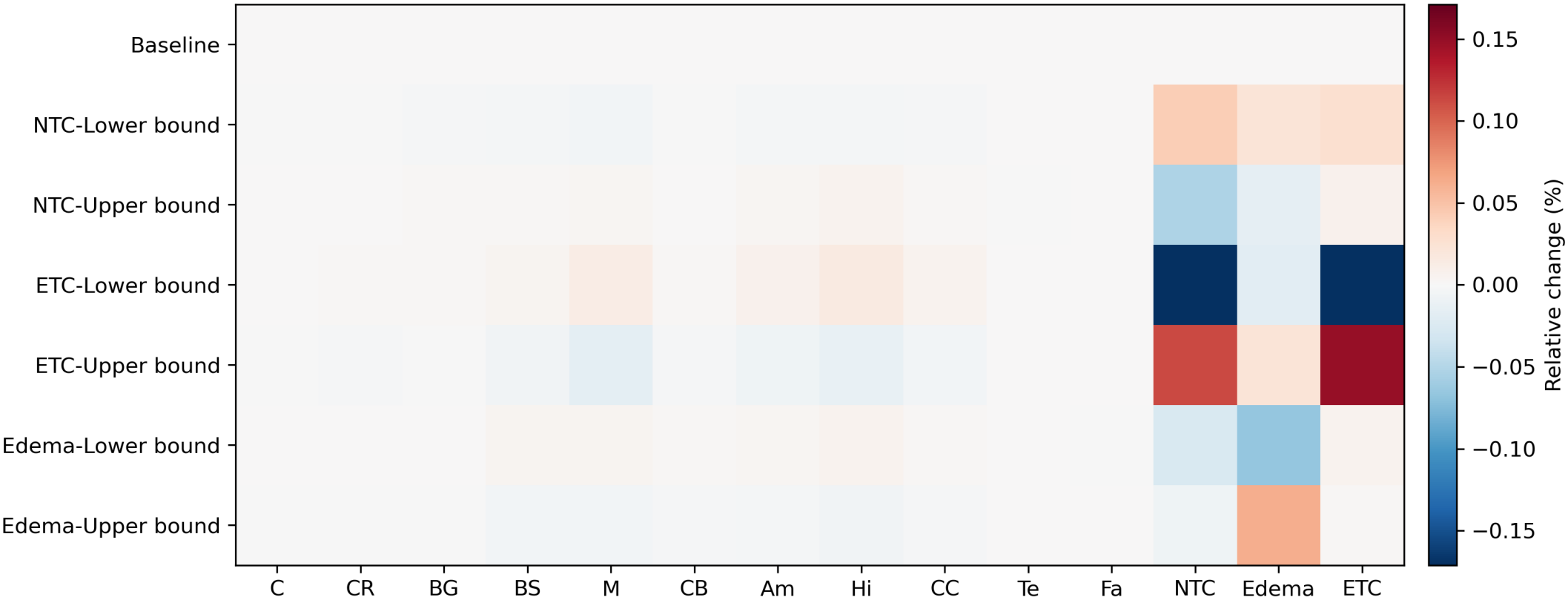
Control case sensitivity of regional third principal stretch to tumor subregion stiffness perturbations. Values indicate percentage change relative to the baseline simulation.

The largest sensitivity is observed for ETC perturbations. Decreasing *µ*_*ETC*_ leads to large reductions in the third principal stretch within both the ETC and NTC regions, with only very small changes observed in the edema region. Inversely, increasing *µ*_*ETC*_ leads to pronounced increases in the third principal stretch within the ETC and NTC regions, again with only weak effects in edema.

This is followed by edema perturbations, where decreasing *µ*_*Edema*_ mainly reduces the third principal stretch within the edema itself, while increasing *µ*_*Edema*_ increases the third principal stretch within the edema region. Compared with ETC, the influence of edema remains more localized and weaker.

NTC perturbations exhibit the opposite trend. Decreasing *µ*_*NTC*_ leads to mild increases in the third principal stretch across all three tumor regions, whereas increasing *µ*_*NTC*_ produces slight reductions in compressive stretch throughout the tumor.

###### Summary

Overall, perturbations of the ETC stiffness produce the largest and most spatially distributed mechanical variations across the brain, dominating the displacement, von Mises stress, and third principal stretch responses. Edema perturbations produce similar but weaker and more diffuse effects, suggesting a secondary buffering role in deformation propagation.

In contrast, perturbations of *µ*_*NTC*_ produce only minor effects on displacement and von Mises stress but generate the strongest sensitivity in the first principal stretch fields, particularly within the NTC itself. Since the NTC does not undergo growth, its stiffness primarily affects localized tensile deformation within the tumor.

Varying the tumor stiffness across its reported range changed the displacement by at most approximately 1.5%, the von Mises stress by at most approximately 4%, and the principal stretches by at most approximately 0.2%. Within these small changes, the ETC stiffness had the largest influence on the global response, while edema acted as a secondary transmission layer and the passive NTC mainly influenced local deformation within the tumor.

#### 3.2.2. Influence of tumor composition on mechanical sensitivity

Three subjects from the BraTS 2021 dataset were selected to assess how tumor composition influences the sensitivity patterns established in the control case, summarized in Table 2. Subject 1 is strongly ETC-dominated with near-absent edema, subject 2 is edema-dominated with minimal NTC, and subject 3 has a composition comparable to the control, but a tumor volume more than 3 times that of the control relative to total brain volume (5.96% vs. 1.71%). The analysis below focuses on displacement magnitude and von Mises stress. Corresponding sensitivity heatmaps for the principal stretch fields are provided in Figures B.13 and B.14 in Appendix B.

**Table 2.** Tumor subregion volume fractions for selected subjects.

| Subject | NTC (%) | ETC (%) | Edema (%) | Tumor/brain (%) |
|---|---|---|---|---|
| Control Subject (01249) | 30.7 | 34.6 | 34.6 | 1.71 |
| Subject 1 (00641) | 25.7 | 74.2 | 0.2 | 2.75 |
| Subject 2 (00636) | 4.7 | 32.4 | 62.9 | 2.81 |
| Subject 3 (00399) | 25.7 | 36.1 | 38.2 | 5.96 |

##### Displacement

Figure 11 shows the percentage change in displacement magnitude for the three selected subjects, computed on a common color scale to allow direct cross-subject comparison. Across all three subjects, the ETC-dominant sensitivity hierarchy established in the control is qualitatively preserved: perturbations of *µ*_*ETC*_ produce the largest and most spatially distributed displacement changes, edema perturbations yield only secondary effects, and NTC perturbations remain localized. The magnitude of the ETC sensitivity differed between subjects. Subject 1, in which ETC comprises 74.2% of the tumor, exhibits the strongest displacement response, with pronounced changes extending across nearly all healthy regions, including the basal ganglia, corpus callosum, cerebellum, tentorium, and falx. By contrast, Subject 2 shows a visibly reduced ETC signal across all healthy regions, with the tentorium and falx, which were among the most sensitive regions in the control, showing considerably weaker responses. Since its ETC fraction (32.4%) is similar to that of the control (34.6%), this attenuation cannot be attributed to ETC fraction, it may instead reflect the large edema shell surrounding the ETC or the tumor location. Subject 3, whose ETC fraction is also comparable to that of the control (36.1%), produces an ETC sensitivity pattern of similar magnitude, with slightly amplified responses in the NTC column consistent with its larger absolute tumor volume. Edema perturbations produce negligible displacement changes across all three subjects, regardless of edema volume fraction. In Subject 2, despite edema comprising 62.9% of the tumor, the edema rows remain essentially flat across all healthy and tumor regions, confirming that the small stiffness contrast between the edema region and surrounding tissue is insufficient to produce measurable displacement sensitivity, even at large volume fractions. In Subject 1, the edema shows very little sensitivity, here due to the low edema volume fraction (0.2%). Together, these subjects provide complementary evidence that edema stiffness does not meaningfully influence the displacement field under the considered parameter ranges. NTC perturbations remain localized across all subjects. Subject 3 shows a comparatively stronger NTC self-response than the control, due to its larger absolute NTC volume rather than a higher NTC fraction. Subject 1 shows moderate NTC sensitivity with a notable response in the brainstem, likely due to the anatomical location of this tumor relative to that structure. Subject 2 shows negligible NTC signal, consistent with its minimal NTC volume fraction (4.7%).

**Figure 11.**
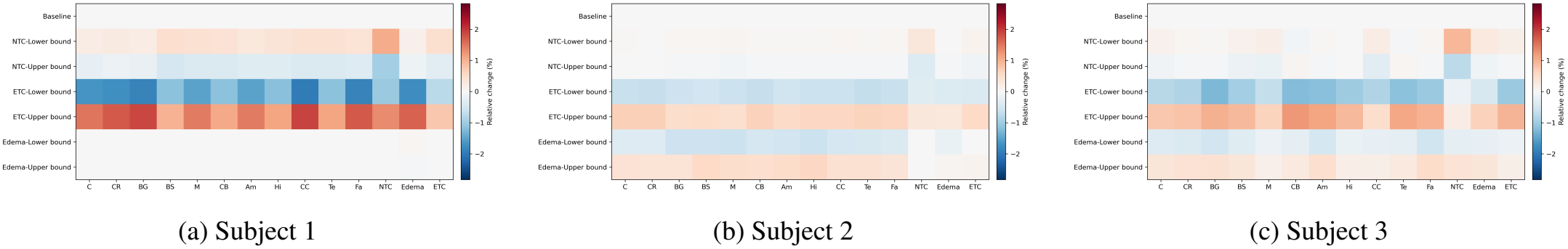
Regional displacement sensitivity for the three representative patient-specific tumors: (a) ETC-dominant tumor, (b) edema-dominant tumor, and (c) large mixed-composition tumor. Values indicate percentage change relative to the baseline simulation and are shown using a common color scale.

##### Von Mises stress

Figure 12 presents the percentage change in regional von Mises stress for the three selected subjects using a common color scale. Across all subjects, the ETC-dominant sensitivity pattern observed in the control subject is preserved, although both the magnitude and spatial distribution of the response vary with tumor composition.

**Figure 12.**
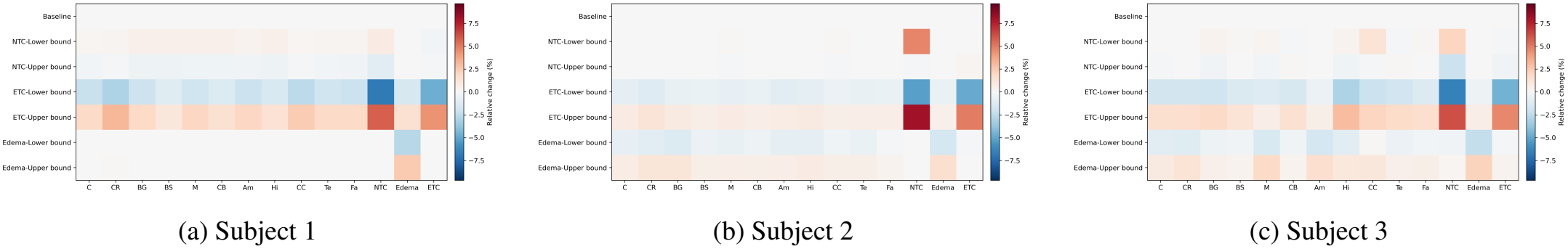
Regional von Mises stress sensitivity for the three representative patient-specific tumors: (a) ETC-dominant tumor, (b) edema-dominant tumor, and (c) large mixed-composition tumor. Values indicate percentage change relative to the baseline simulation and are shown using a common color scale.

For all three brains, perturbations in ETC stiffness produce stress changes across healthy regions that vary in magnitude with tumor composition. The strongest sensitivities remain restricted within the tumor subregions. Among the three subjects, Subject 1 exhibits the strongest stress sensitivity within the surrounding healthy tissue, which is consistent with its high ETC volume fraction, where ETC constitutes approximately 74% of the total tumor volume. As in the control subject, the NTC and ETC regions show the most pronounced stress responses under ETC perturbations, whereas the edema region consistently exhibits weak sensitivity.

Similarly, perturbations in edema stiffness produce only minor stress changes across all regions in all three subjects. This trend remains evident even in Subject 2, where edema constitutes 62.9% of the total tumor volume, with only weak changes observed locally within the edema region itself, with the sign following the direction of the stiffness perturbation. These results suggest that the mechanical contrast associated with edema is too small to generate substantial von Mises stress sensitivity, regardless of its volume fraction within the tumor.

A distinct behavior is observed for NTC perturbations. In Subject 2, a clear and localized stress response develops within the NTC region despite its very small volume fraction (4.7%). In contrast, the response is weaker in subjects 1 and 3, where the NTC takes up approximately 25% of the tumor volume. This behavior is likely related to the passive mechanical role of the NTC: its mechanical response is primarily influenced by compression from the surrounding growing regions. In Subject 2, the small NTC core is surrounded by a large volume of actively growing ETC and edema tissue, which together account for 95.3% of the tumor volume. The resulting confinement generates a relatively high local stress concentration within the NTC. In subjects 1 and 3, the larger NTC region distributes this compressive loading over a greater volume, leading to lower stress intensity.

Overall, the von Mises stress analysis confirms and extends the displacement field observations: ETC stiffness remains the dominant driver of global stress sensitivity, its influence on surrounding healthy tissue increases with ETC volume fraction, and edema stiffness continues to play a mechanically secondary role even at high volume fractions. Additionally, the von Mises stress analysis reveals that NTC stress sensitivity is governed not by its own volume fraction alone, but by the relative volume of surrounding growing tissue, an effect that amplifies internal stress redistribution when the passive core is small relative to the actively expanding tumor mass.

##### Summary

The control case established the intrinsic mechanical role of each tumor subregion independently of compositional variability: ETC stiffness dominated the global mechanical response, edema produced weaker and more diffuse effects, and NTC stiffness primarily influenced localized tensile deformation within the tumor. The patient-specific analyses showed that these sensitivity hierarchies remain qualitatively preserved across heterogeneous tumors, but that their magnitude and spatial extent depend strongly on tumor composition. In particular, ETC-dominant tumors amplified displacement and stress propagation into surrounding brain tissue, whereas edema-dominant tumors remained mechanically less sensitive despite large edema volumes. Overall, these results indicate that, within the reported stiffness ranges, uncertainty in tumor material properties has only a small effect on the predicted mass effect compared with other modeling choices, such as the growth law or the dural structures. Among the tumor subregions, the ETC had the largest influence.

## 4. Conclusion

This study investigated the mechanical role of intracranial dural structures and tumor subregion material properties in finite element models of brain tumor mass effect. It evaluated the influence of the falx cerebri and tentorium cerebelli on deformation and stress redistribution, and assessed how tumor-subregion stiffness affects the simulated local and global mechanical response of the brain.

A heterogeneous nine-region finite element brain model was generated from MRI data and included the falx cerebri, tentorium cerebelli, and a tumor. Two analyses were performed. First, the mechanical influence of the dural structures was assessed by comparing models with and without the falx and tentorium using a synthetic lesion configuration. Second, a parametric study was conducted using patient-specific tumor models to evaluate how variations in the stiffness of the necrotic core, enhancing tumor core, and edema affect displacement, stress, and stretch fields.

The dura mater analysis showed that the falx cerebri and tentorium cerebelli had only a minor influence on peak displacement, but a larger influence on the von Mises stress field. Their inclusion therefore matters little for estimating the global magnitude of mass effect, but noticeably alters the spatial distribution of displacement and stress, particularly near the interhemispheric region and the tentorial interface. The dura acted as a stiff internal load-bearing membrane that mechanically separated supratentorial and infratentorial regions and limited the transmission of deformation across these partitions. These findings indicate that dural structures should be considered when studying directional deformation and stress redistribution.

This is further supported by the analysis of ventricular volume under sustained tumor growth. With the falx present, both lateral ventricles were compressed as the tumor grew, and the total ventricular volume decreased monotonically. Without it, the ventricle on the tumor side expanded while the ventricle in the opposite hemisphere was compressed, resulting in a partial recovery of the total ventricular volume. The falx thus acts as an internal wall that limits the displacement of the ventricle in the hemisphere containing the tumor and transmits the tumorinduced load across the midline.

The tumor material parametric analysis showed that varying the tumor subregion stiffness across its reported range changed displacement and stress by only a few percent. Among the subregions, variations in ETC stiffness produced the largest changes, particularly in tumors with large ETC volume fractions. Edema stiffness had a weaker and more diffuse influence, acting mainly as a buffering layer. Although edema can occupy a large fraction of the tumor-associated volume, its stiffness remained close to that of healthy white matter within the physiological ranges considered, resulting in relatively small sensitivity effects. In contrast, NTC stiffness mainly affected the principal stretch within the tumor itself, with minimal influence on displacement or stress propagation within the brain.

Overall, the study demonstrates that the explicit representation of intracranial structures alters the simulated mechanical response in brain tumor mass-effect simulations. The results further suggest that the global mechanical behavior of growing brain tumors is primarily governed by the actively growing ETC tissue, whereas edema and necrotic regions mainly influence local deformation patterns and stress redistribution.

These findings should, however, be interpreted strictly as mechanistic observations within the assumptions of the present finite element framework. The study was designed to isolate and characterize the relative mechanical influence of dural structures and tumor subregion properties, rather than to establish a clinically predictive model of tumor-induced brain deformation. The reported displacement, stress, stretch, and ventricular-volume responses must be validated against longitudinal patient data, intraoperative measurements, or independent experimental observations.

Several limitations should be acknowledged. First, tumor growth was modeled using a prescribed isotropic expansion rather than a coupled growth model that incorporates cell proliferation. Second, the constitutive formulation was purely hyperelastic, omitting the poroelastic or viscoelastic effects that can influence edema evolution and long-term tissue response. Furthermore, the patient-specific geometries were reconstructed from preoperative MRI in which the tumor and its mass effect were already present. These configurations were treated as stress-free reference states, and growth was prescribed on top of them. Consequently, residual stresses and deformations accumulated during prior tumor growth were neglected. In addition, the dural structures were tied to the adjacent brain tissue, whereas *in vivo* some relative sliding is possible. Moreover, the parameters of the tumor material were derived from MRE-reported stiffness ranges obtained under small-deformation conditions, rather than from large-strain mechanical measurements. Finally, because the study relied on a one-at-a-time parameter variation method, it did not evaluate the potential interaction effects between the material parameters of the tumor subregions.

Future work should include coupling biomechanical growth with reaction–diffusion tumor models, and incorporate longitudinal MRI data for validation of predicted deformation and stress fields. Further studies should also quantify how uncertainty in tumor segmentation, boundary conditions, and regional brain material properties affects predicted mass effect. Such developments will be important for translating patient-specific finite element brain tumor models into reliable tools for prediction of tumor progression, surgical planning, and treatment simulation.

## Acknowledgements

We acknowledge the financial support by the Deutsche Forschungsgemeinschaft (DFG, German Research Foundation) through the grant BU 3728/1-1. Additionally, this result is part of a project that has received funding from the European Research Council (ERC) under the European Union’s Horizon 2020 research and innovation programme (Grant agreement No. 101116420-MAGERY).

## Data availability

Data will be made available upon reasonable request.

## Appendix A. Material parameters

**Table A.3.** Different brain regions and their *α* and *µ* values according to experiments from [38].

| Brain region | Preconditioned<br>ν = 0.45 |  | Preconditioned<br>ν = 0.49 |  | Unconditioned<br>ν = 0.45 |  | Unconditioned<br>ν = 0.49 |  |
|---|---|---|---|---|---|---|---|---|
|  | α | μ | α | μ | α | μ | α | μ |
| Amygdala | -19.67 | 257.05 | -14.16 | 193.34 | -14.74 | 386.97 | -10.10 | 293.06 |
| Basal ganglia | -18.95 | 252.15 | -15.05 | 169.67 | -15.90 | 338.67 | -11.49 | 246.21 |
| Brain stem | -20.41 | 233.12 | -19.59 | 118.06 | -16.82 | 336.44 | -13.60 | 225.08 |
| Cortex | -19.27 | 281.75 | -16.72 | 175.21 | -16.16 | 392.24 | -12.16 | 285.97 |
| Cerebellum | -23.22 | 177.49 | -18.79 | 115.36 | -19.49 | 255.38 | -14.08 | 196.55 |
| Corpus callosum | -14.94 | 61.66 | -13.72 | 37.64 | -12.82 | 95.43 | -9.11 | 67.82 |
| Corona radiata | -19.70 | 157.09 | -19.08 | 87.58 | -16.93 | 218.75 | -13.55 | 146.54 |
| Hippocampus | -17.37 | 184.21 | -13.09 | 129.96 | -13.25 | 292.58 | -9.23 | 218.74 |
| Midbrain | -18.68 | 215.70 | -15.15 | 137.67 | -15.11 | 314.92 | -10.89 | 227.69 |

**Table A.4.** Mesh size of the finite element models.

| Model | Elements | DOFs |
|---|---|---|
| Synthetic lesion | 175,145 | 561,150 |
| Control (01249) | 245,768 | 781,848 |
| 00641 (1) | 267,542 | 849,717 |
| 00636 (2) | 244,192 | 776,805 |
| 00399 (3) | 232,267 | 739,782 |

## Appendix B. Sensitivity heatmaps

**Figure B.13.**
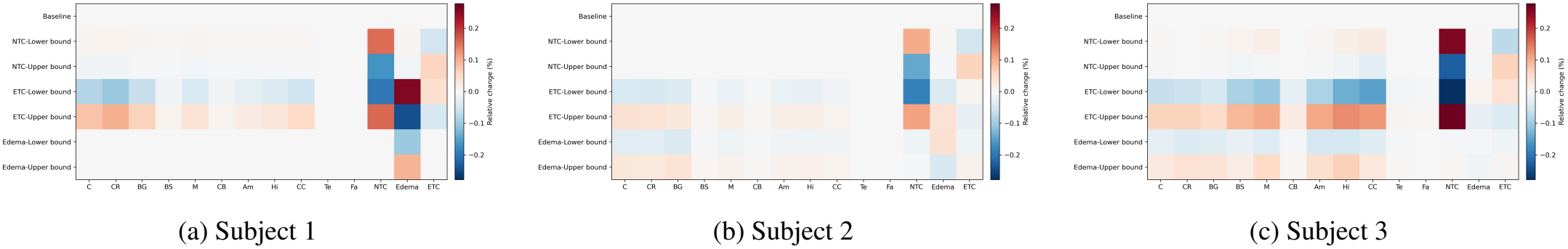
Regional first principal stretch sensitivity for the three representative patient-specific tumors: (a) ETC-dominant tumor, (b) edema-dominant tumor, and (c) large mixed-composition tumor. Values indicate percentage change relative to the baseline simulation.

**Figure B.14.**
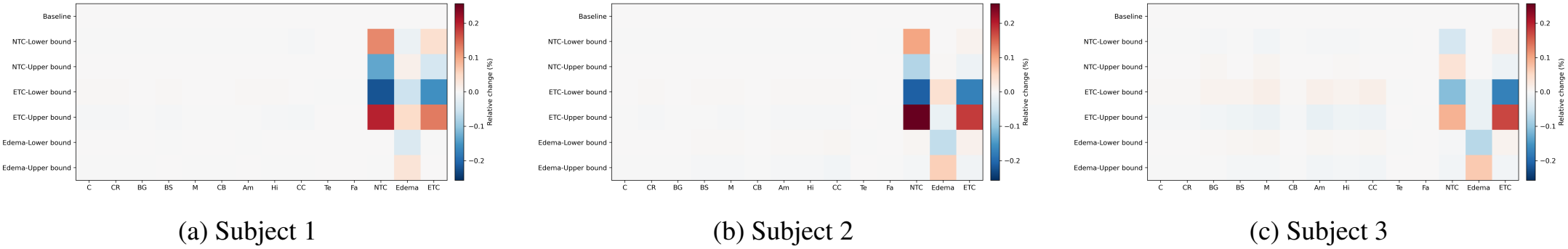
Regional third principal stretch sensitivity for the three representative patient-specific tumors: (a) ETC-dominant tumor, (b) edema-dominant tumor, and (c) large mixed-composition tumor. Values indicate percentage change relative to the baseline simulation.

## Appendix C. Supratentorial synthetic tumor

**Figure C.15.**
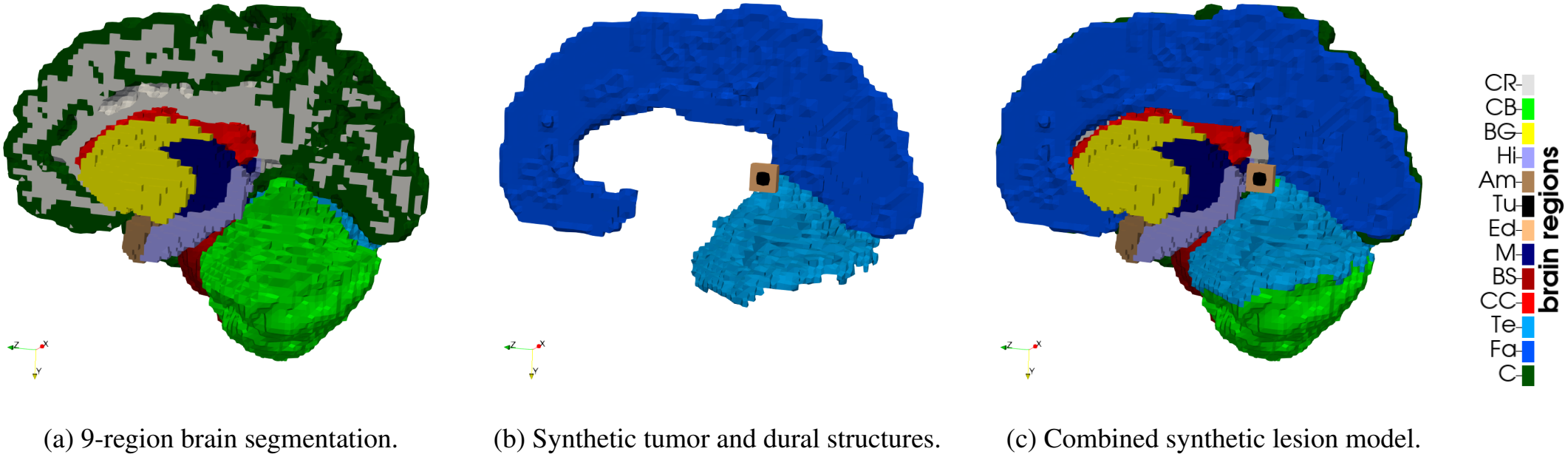
Finite element model used for the synthetic lesion study. The model includes nine mechanically distinct brain regions, dural structures represented by the falx cerebri and tentorium cerebelli, and a supratentorial synthetic tumor with surrounding edema.

**Figure C.16.**
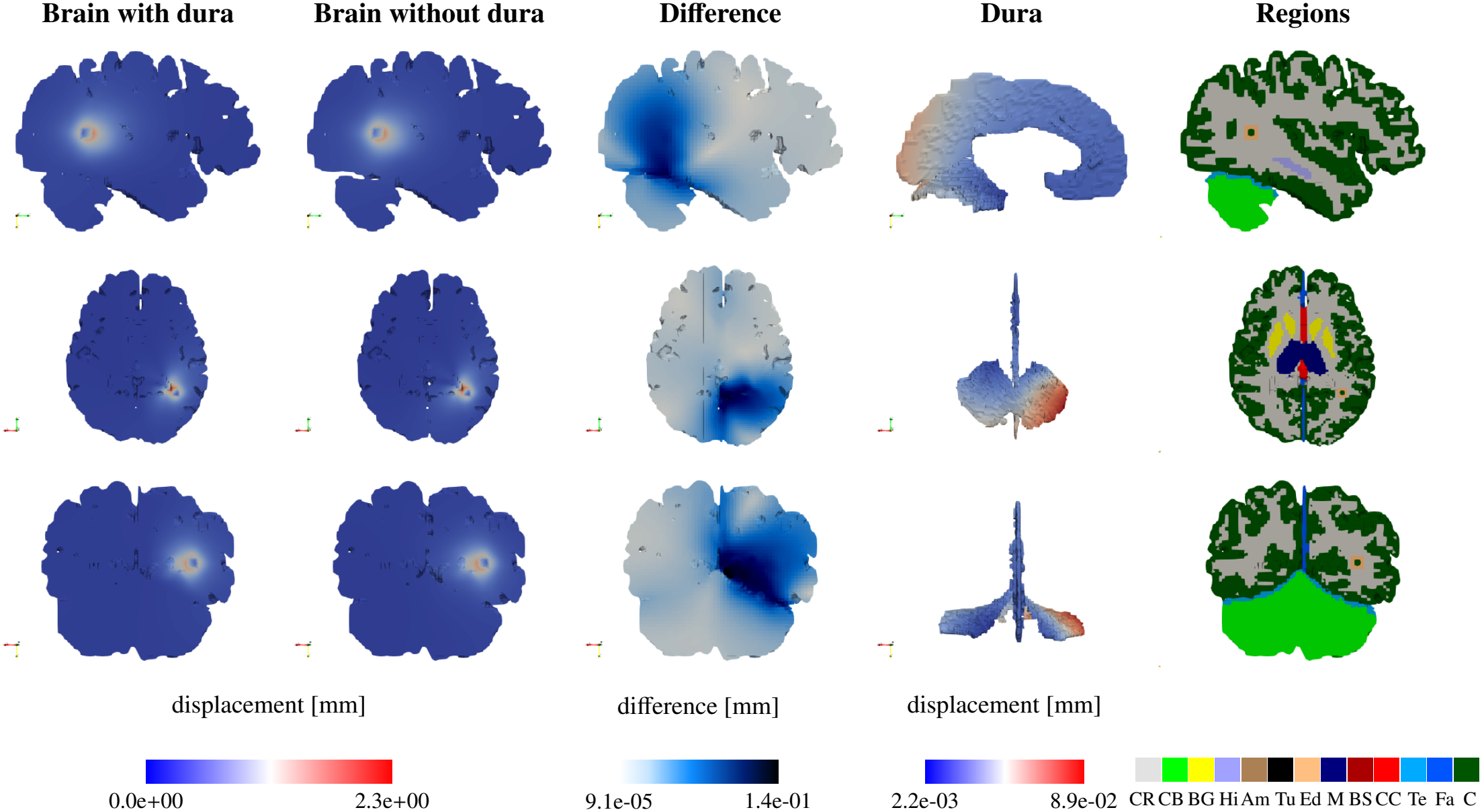
Displacement field [mm] comparison between models with and without dural structures for the synthetic lesion case. Columns show displacement magnitude with dura, displacement magnitude without dura, absolute displacement difference, displacement of the dural structures, and anatomical regions. Rows correspond to sagittal, axial, and coronal sections.

**Figure C.17.**
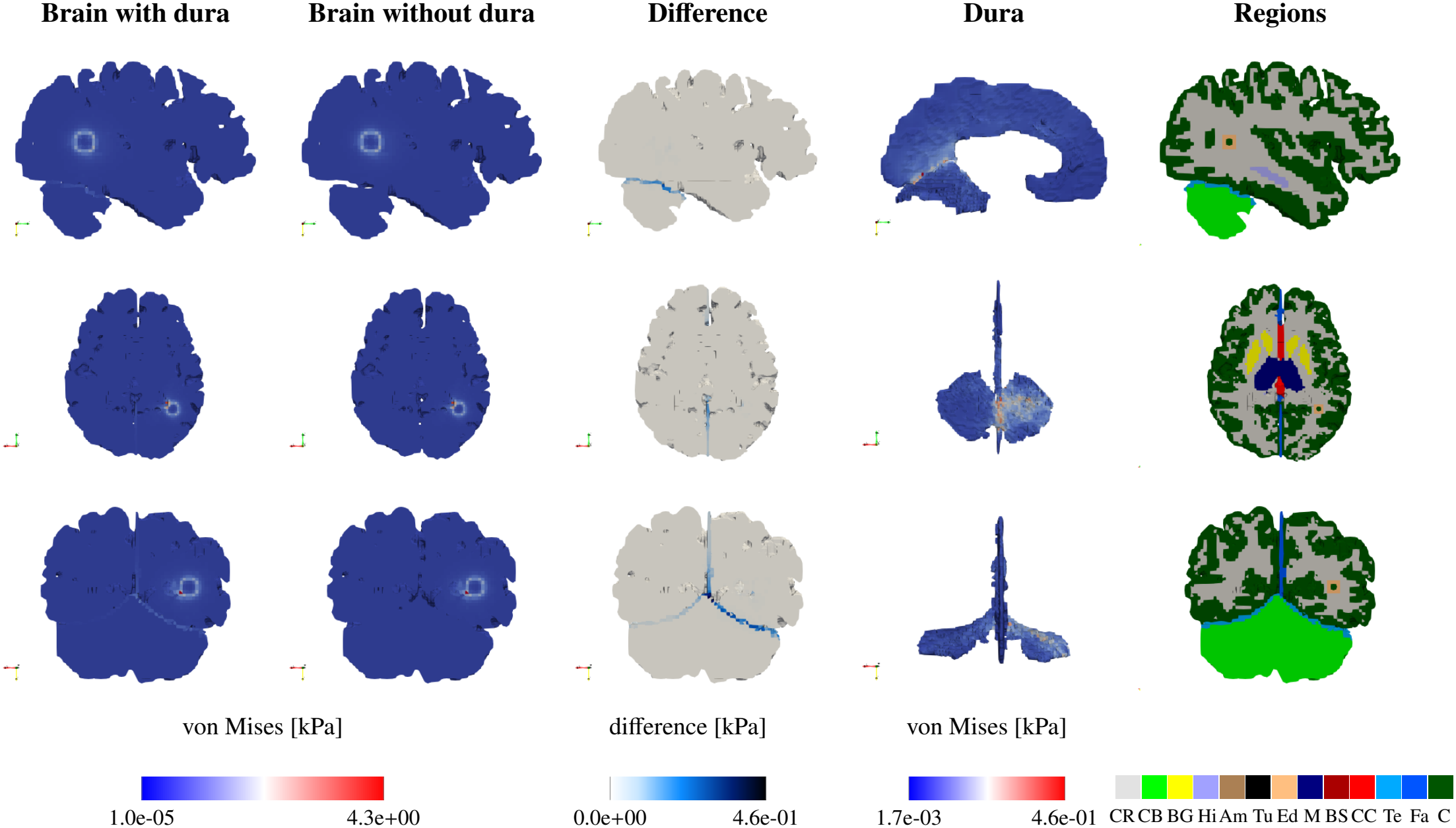
Von Mises stress [kPa] comparison between models with and without explicit dural structures for the synthetic lesion case. Columns show von Mises stress with dura, von Mises stress without dura, absolute stress difference, stress carried by the dural structures, and anatomical regions. Rows correspond to sagittal, axial, and coronal sections.

## Footnotes

1 https://sites.wustl.edu/oasisbrains/

2 https://www.kaggle.com/datasets/dschettler8845/brats-2021-task1

3 https://github.com/BRAINIACS-Group/Brain-Creation-Code

4 https://github.com/BRAINIACS-Group/EFI_atrophy_and_tumor

